# PIP_2_ promotes conformation-specific dimerization of the EphA2 membrane region

**DOI:** 10.1101/2020.10.14.338293

**Authors:** Katherine M. Stefanski, Charles M. Russell, Justin M. Westerfield, Rajan Lamichhane, Francisco N. Barrera

## Abstract

The impact of the EphA2 receptor on cancer malignancy hinges on the two different ways it can be activated. EphA2 induces anti-oncogenic signaling after ligand binding, but ligand-independent activation of EphA2 is pro-oncogenic. It is believed that the transmembrane (TM) domain of EphA2 adopts two alternate conformations in the ligand-dependent and the ligand-independent states. However, it is poorly understood how the difference in TM helical crossing angles found in the two conformations impacts the activity and regulation of EphA2. We devised a method that uses hydrophobic matching to stabilize two conformations of a peptide comprising the EphA2 TM domain and a portion of the intracellular juxtamembrane (JM) segment. The two conformations exhibit different TM crossing angles, resembling the ligand-dependent and ligand-independent states. We developed a single-molecule technique using SMALPs to measure dimerization in membranes. We observed that the signaling lipid PIP_2_ promotes TM dimerization, but only in the small crossing angle state, which we propose corresponds to the ligand-independent conformation. In this state the two TM are almost parallel, and the positively charged JM segments are expected to be close to each other, causing electrostatic repulsion. The mechanism PIP_2_ uses to promote dimerization might involve alleviating this repulsion due to its high density of negative charges. Our data reveal a conformational coupling between the TM and JM regions, and suggest that PIP_2_ might directly exert a regulatory effect on EphA2 activation in cells that is specific to the ligand-independent conformation of the receptor.

## Introduction

Eph receptor tyrosine kinases (RTKs) are the largest family of RTKs in humans. Eph receptors are involved in tissue patterning during embryonic development, neuronal plasticity, and wound healing (1, 2). Beyond their normal physiological functions, Eph receptors can contribute to human diseases. For example, elevated EphA4 signaling results in neuronal damage in Alzheimer’s disease and amyotrophic lateral sclerosis (ALS) (3–6), and the loss of EphB2/B3 signaling is implicated in skeletal malformations that cause cleft palate (7). Moreover, a large body of research exists establishing that Eph receptors are overexpressed in a variety of cancer types. Specifically, EphA2 overexpression is found in breast, ovarian, prostate, and pancreatic cancers and is correlated with aggressive tumors, high rates of tumor recurrence, and poor patient prognosis (8–14). Additionally, Eph receptors have been found to be cellular receptors for viruses that cause cancer. For example, EphA2 is a receptor for Kaposi’s sarcoma-associated herpes virus and Epstein-Barr virus (15, 16).

EphA2 signaling pathways control cell proliferation, migration, and cell retraction (17). EphA2 can engage in two modes of activation: ligand-dependent and ligand-independent (i.e., non-canonical). Ligand-dependent signaling requires activation of EphA2 by binding of its ligand, ephrinA1, resulting in the phosphorylation of residues Y588, Y594 and Y772. This results in signaling that inhibits metastatic phenotypes by causing cell retraction/rounding and decreasing cell proliferation and migration (1, 18, 19). Conversely, ligand-independent signaling is responsible for oncogenic phenotypes and occurs via phosphorylation of S897 by the kinases AKT, RSK, or PKA (18, 20–22). Overexpression of EphA2 in cancers is often accompanied by a loss of ephrinA1 ligands (23, 24). It is believed that this imbalance of EphA2 and ephrinA1 results in both increased ligand-independent signaling and a decreased ligand-dependent signaling, promoting tumor growth and malignancy (25).

Due to its prominent role in oncogenesis, EphA2 has become an attractive drug target and, as such, an active area of research. The structure of EphA2 includes an extracellular ligand-binding domain, a single-pass transmembrane domain (TMD), and an intracellular kinase domain connected to the TMD by a disordered juxtamembrane (JM) segment. In the ligand-independent state, EphA2 exists in a monomer-dimer equilibrium (26). Upon binding of ephrinA1, EphA2 dimerization via the ligand-binding domains is promoted and leads to the formation of large signaling clusters (27). While some aspects of Eph receptor signaling have been elucidated, the exact molecular events of how the receptor transmits an extracellular signal across the plasma membrane remain unknown. As a membrane-spanning receptor, the TMD is expected to play a role in conferring signals across the plasma membrane.

An NMR structure of the EphA2 TMD dimer was solved by Bocharov *et al*. (28). This dimer had a small interhelical crossing angle (15°), and the interface was mediated by a heptad repeat (HR) motif. The same study conducted molecular dynamics (MD) simulations on the NMR structure and found that the EphA2 TMD dimer could rotate to a glycine zipper (GZ) interface with a larger interhelical crossing angle (45°). It was hypothesized that the receptor switches between these two conformations with different dimerization interfaces. In a follow-up study, it was reported that mutations in the HR motif decreased ligand-independent signaling, while mutations in the glycine zipper decreased ligand-dependent signaling (29). These findings, combined with the structural data, gave rise to the following model: In the ligand-independent signaling state the EphA2 TMD dimerizes via the HR motif with nearly parallel TMDs, while in the ligand-dependent signaling state, EphA2 dimerizes via the GZ with tilted TMDs (28, 29). It is unknown how the switch in dimerization interface and opening of the interhelical crossing angle participate in conferring the extracellular signal from the outside of the cell to the cytoplasm. It is further unknown if the JM responds to changes in the TMD.

JM-lipid interactions are involved in the activation of several receptors. These interactions are believed to be mediated by positively charged JM residues interacting with negatively charged lipid head groups. For EphA2, associations between basic JM residues and phosphatidylinositol 4,5-bisphosphate (PIP_2_) have been computationally predicted (30). Notably, the first five positively charged residues (HRRRK) of the EphA2 JM are predicted to establish strong interactions with negatively charged PIP_2_. However, this association has not been experimentally observed. Interactions between the JM and PIP_2_ might regulate the activity of EphA2, since the two signaling modalities of EphA2 alter PIP_2_ levels. Ligand-dependent signaling activates PI3K, which phosphorylates PIP_2_ to generate PIP_3_ (1). While ligand-independent signaling recruits SHIP2, which converts PIP_3_ to PIP_2_ and can be triggered by AKT upon PI3K activation (17). However, it is unknown if these local changes in PIP_2_ directly alter EphA2 signaling.

In the present study, we investigate how the TMD and JM regions of EphA2 are affected by signaling-related changes in TMD orientation and lipid environment. We use hydrophobic matching to stabilize the two conformations of the EphA2 TMD region. We also examine how bilayer composition and the position of JM residues affects self-assembly of the TMD of EphA2, using a novel single-molecule fluorescence approach in styrene maleic acid lipid particles (SMALPs). Our results indicate that PIP_2_ specifically promotes dimerization of one of the two TMD conformations via interactions with the JM. Implications for the role of lipids in the two signaling states of EphA2 are discussed.

## Results

### Bilayer thickness drives changes in TM orientation

We sought to generate an *in vitro* model of the ligand-independent and ligand-dependent signaling states of the membrane region of the EphA2. To this end, we used the TMJM peptide, which comprises a short stretch of extracellular residues, the TMD and first 5 JM residues of EphA2 (Fig. 1A). At the C-terminus, we added a cysteine to enable dye conjugation, and a tryptophan as a fluorescent reporter of the JM segment. As noted above, the two EphA2 signaling configurations have different inter-helical crossing angles. To promote the different configurations, we used model membranes composed of 14:1 PC or 22:1 PC, which only differ in the length of their acyl chains. Since 22:1 PC contains 8 more tail carbons than 14:1 PC, it forms thicker bilayers (45.5 Å vs. 29.6 Å) (31). Our hypothesis was that we could use hydrophobic matching to stabilize the TMD of EphA2 in the two different conformations (32). Specifically, TMJM would orient closer to the bilayer normal in a thick bilayer (22:1 PC), while it would tilt further away from the bilayer normal in a thin (14:1 PC) bilayer. Thus, we sought to use bilayer composition to tune helical tilt and recapitulate the ligand-independent and ligand-dependent signaling configurations.

**Figure 1.**
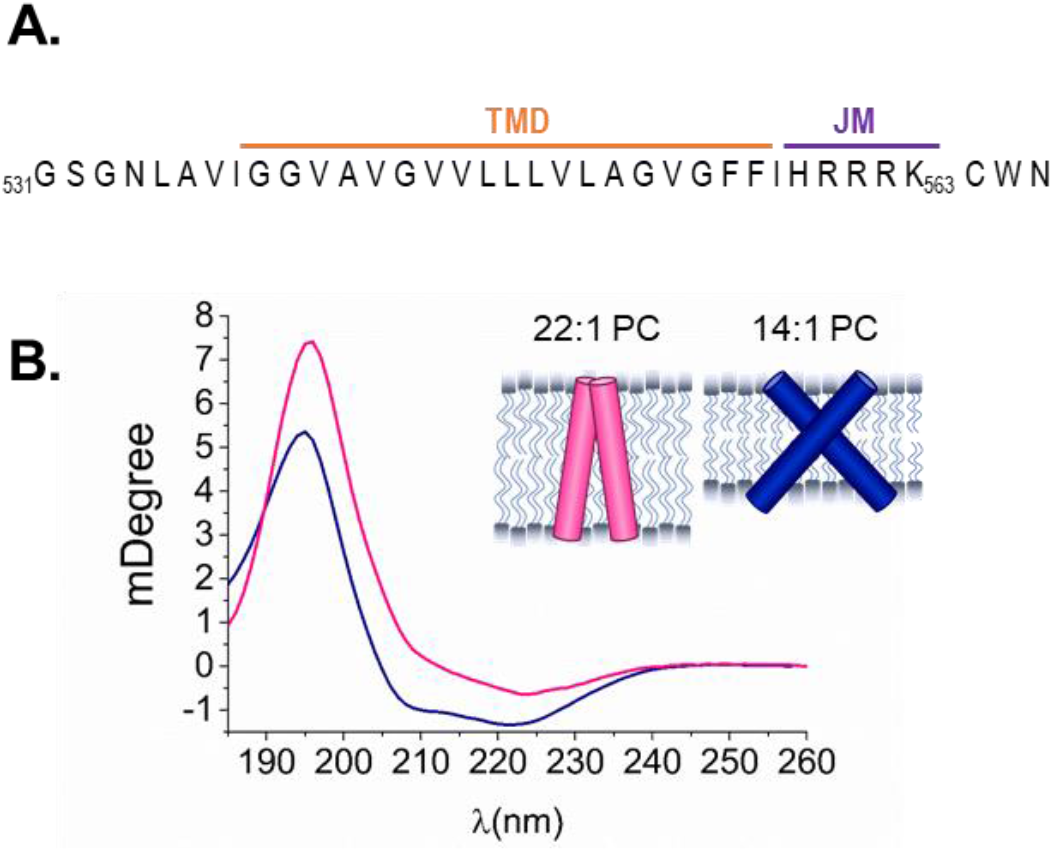
Bilayer thickness drives differences in TMJM helical tilt. **A.** Sequence of the TMJM peptide comprised of EphA2 residues 531-563 with added CWN residues at the C-terminus. **B.** OCD spectra of TMJM in 22:1 PC (fuchsia) and 14:1 PC (navy). Inset: model of the conformations of TMJM in 14:1 PC and 22:1 based on OCD data.

We used oriented circular dichroism (OCD) to test the effects of bilayer thickness on the helical tilt of TMJM, when reconstituted in supported lipid bilayers composed of 14:1 PC or 22:1 PC. Fig. 1B shows that the obtained OCD spectra had two α-helical minima at 208 and 224 nm, indicating a transmembrane orientation. To assess the helical tilt, we can examine changes in the 208 nm minimum. A more positive value indicates a less tilted helix, as seen in 22:1 PC, while a lower value indicates a more tilted helix, as seen in 14:1 PC (Fig. 1B) (33–35). These data suggest that, on average, the TMDs are more vertical in thicker bilayers, like the ligand-independent signaling configuration. In contrast, the TMDs are more tilted in thinner bilayers, which might correspond to the ligand-dependent signaling configuration. The observed changes in tilt were independent of lipid to peptide ratio and were also observed at lower peptide concentrations (Fig. S1). Standard circular dichroism was performed on TMJM in 14:1 PC and 22:1 PC vesicles to ensure that the secondary structure of the peptide was similar in both lipids (Fig. S1). The OCD results suggest that the peptide responds to being placed in bilayers of different thickness by adjusting helical tilt, providing a means to further study the two observed conformations.

### TMJM can form dimers in thin and thick bilayers

We wanted to investigate if TMJM self-assembled into biologically relevant dimers. To this end, we developed a single-molecule photobleaching method using SMALPs. TMJM labeled with the fluorescent dye Alexa Fluor 488 was reconstituted in multilamellar vesicles (MLVs) composed of either 14:1 PC or 22:1 PC containing 3% biotinylated phosphatidylethanolamine (biotin-PE). The MLVs were then incubated with styrene maleic acid (SMA). SMA forms a polymer belt around units of lipid bilayer and proteins (Fig. 2A) (36, 37). Transmission electron microscopy (TEM) confirmed that lipid composition did not alter SMALP size (Fig. S2), and co-localization experiments confirmed SMALP composition (Fig. S3). SMALPs were immobilized on a microscope slide via a biotin-streptavidin linkage (Fig. 2A, Fig. S3). Imaging was conducted using total internal reflection fluorescence (TIRF) microscopy. By analyzing the fluorescence of individual SMALPs over time, we can count single fluorophores by detection of photobleaching events (Fig. 2B) to determine the number of labeled TMJM peptides contained in a single SMALP. We found a substantial fraction of the peptide existed in SMALPs yielding two photobleaching steps (Fig. 2C). The majority of SMALPs contained one or two labeled TMJM peptides. Only a small fraction of SMALPs had 3 or more photobleaching steps (Fig. S4). These results were robust as similar values were found for 2 bleaching steps in 22:1 PC and 14:1 PC (Fig. 2C and S1). The single-molecule results suggest that TMJM is in a similar monomer-dimer equilibrium in 14:1 PC and 22:1 PC.

**Figure 2.**
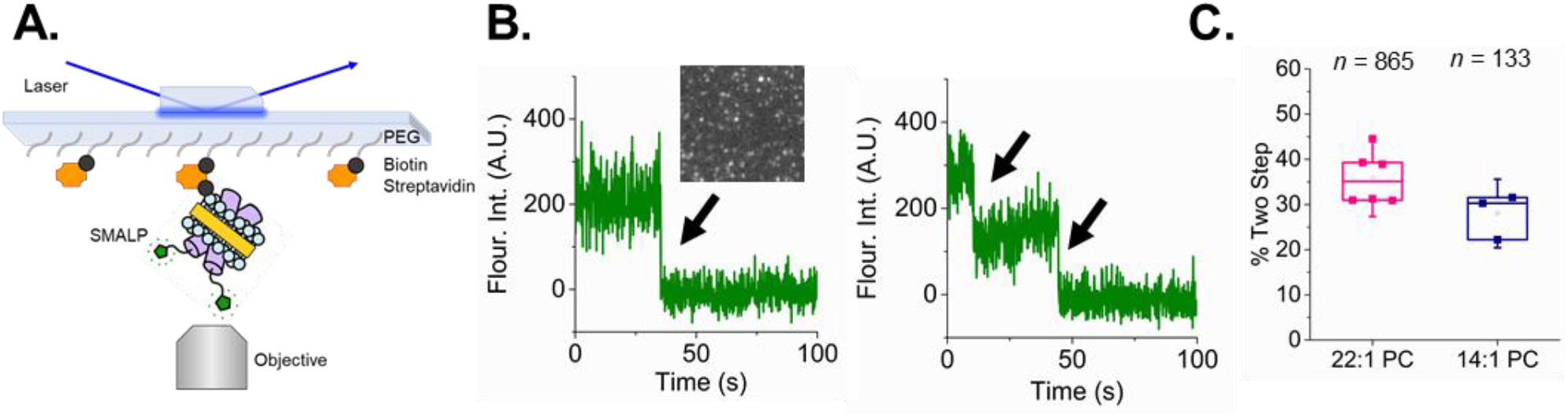
TMJM dimerization observed by single-molecule TIRF of SMALPs. **A.** Schematic of TIRF experimental setup. SMALP showing SMA polymer (yellow) encircling lipids (blue) containing TMJM peptides (purple) labeled with Alexa Fluor 488 (green), immobilized on a PEGylated slide via a biotin (black) and streptavidin (orange) linkage. **B.** Representative fluorescence traces showing one (left) and two (right) photobleaching steps (black arrows). Representative TIRF image of SMALPs (inset). **C.** Box and whiskers plot (upper quartile, median, and lower quartile) showing percentages of peptide counted in traces with two photobleaching steps in 22:1 PC and 14:1 PC at a lipid to peptide ratio of 300:1. Data are from 3-6 independent experiments ± S.D, *n* is total number of traces counted with two photobleaching steps.

### Helical tilt alters the environment of JM residues but not the distance from the bilayer surface

We next investigated if JM-lipid interactions are different in the two conformations adopted by TMJM. We also sought to understand if the change in helical tilt observed for TMJM in thin and thick bilayers altered the position of the JM residues. We used the tryptophan placed after the JM residues as a reporter (Fig. 1A), and fluorescence experiments were performed in 14:1 PC and 22:1 PC vesicles. Sample tubes were washed with SDS and CD was performed to ensure equivalent amounts of the peptide were recovered in each lipid when liposomes were resuspended thus rendering direct comparisons of fluorescence intensities valid (Fig. S8 A). We observed that the fluorescence intensity of the tryptophan was higher in 14:1 PC than 22:1 PC (Fig. 3A) but the spectral maximum was similar (Fig. S5). These data suggest a small change in the environment of the tryptophan that is not related to differences in membrane burial of the JM but may be due to differences in relative orientation (38).

**Figure 3.**
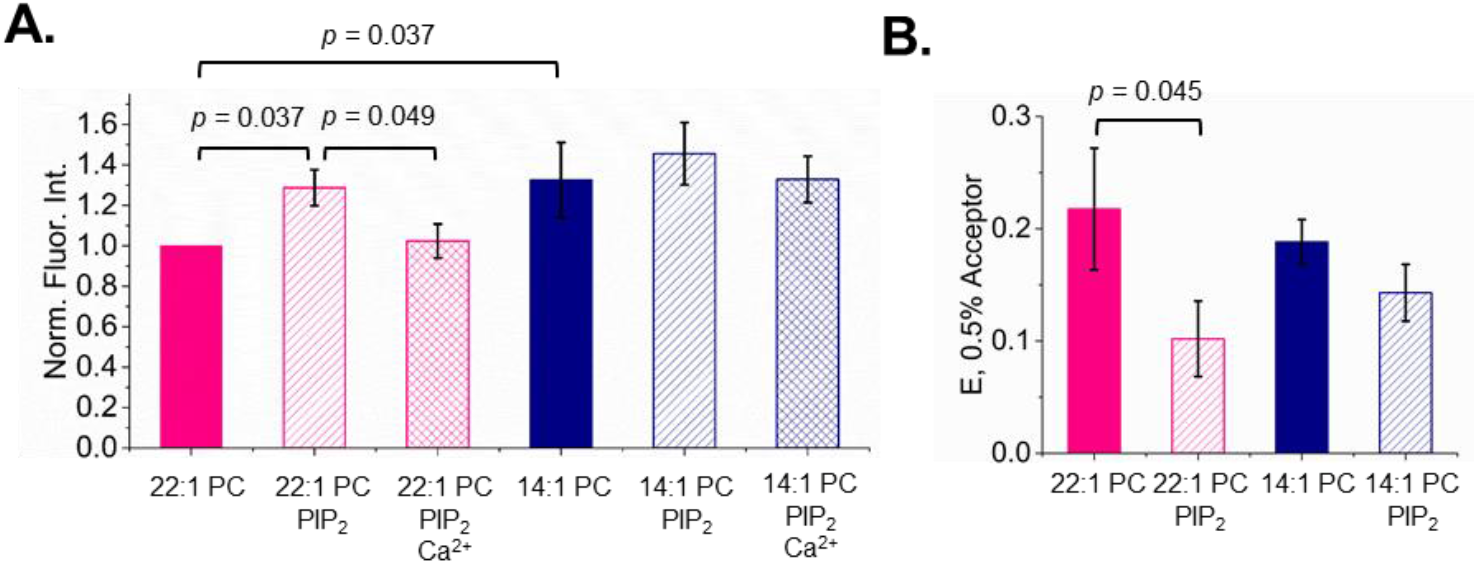
Bilayer thickness and PIP_2_ alter Trp environment while PIP_2_ changes headgroup distance. **A.** Normalized intensities of TMJM Trp in 22:1 PC and 14:1 PC liposomes in the presence and absence of PIP_2_ and 5 mM Ca^2+^. *p*-values were determined by Mann-Whitney U tests, bars are means ± S.D., *n* = 3 **B.** FRET efficiencies calculated from FRET experiments with TMJM (Trp, donor) in 14:1 PC and 22:1 PC liposomes in the presence and absence of 3% PIP_2_ (3% DNS-PE, acceptor) in liposomes. Bars are means ± S.D., *n* = 3). (*p*-value is from one-way ANOVA followed by Tukey post-hoc test.

We next examined the association of the JM with the bilayer interface using the tryptophan as a FRET (Förster Resonance Energy Transfer) donor and a headgroup-labeled dansyl phosphatidylethanolamine (DNS-PE) as an acceptor. In both 14:1 PC and 22:1 PC liposomes, a decrease in donor fluorescence was observed upon the addition of 0.25 - 3% acceptor, indicating that FRET occurred in all conditions (Fig. S6). By calculating the FRET efficiency (E) at 0.5% acceptor we were able to quantify that the FRET occurring in 22:1 PC and 14:1 PC were similar (Fig. 3B). These data, combined with the tryptophan fluorescence results (Fig. 3A), led us to conclude that interhelical crossing angle alone does not significantly affect the association of the JM residues of TMJM with PC lipid bilayers.

### PIP_2_ drives changes in JM-headgroup distance only in thick membranes

To begin examining the effects anionic lipids have on the TMD orientation and JM-lipid associations; we repeated the OCD experiments in the presence of 3% PIP_2_. We observed no large shifts in helical tilt caused by the addition of PIP_2_ in either thin or thick bilayers (Fig. S7). Therefore, we conclude that the addition of anionic lipids does not perturb the hydrophobic matching-driven changes in helical tilt observed in the original OCD experiments (Fig. 1B)

We next determined if PIP_2_ could cause changes in JM-membrane interactions. We performed both tryptophan fluorescence and FRET measurements. When examining tryptophan fluorescence, we added the divalent cation Ca^2+^ as a control to shield the negative charges on PIP_2_. Saturating levels of Ca^2+^ were used in these experiments as determined by calcium influx assays (Fig. S8). In 22:1 PC, we observed a statistically significant increase in tryptophan fluorescence in the presence of PIP_2_ (Fig.3A). The observed intensity increase was reversed upon the addition of Ca^2+^ (Fig. 3A). This result suggests that cationic JM residues participate in electrostatic interactions with the anionic PIP_2_ headgroups and that this interaction is placing the JM tryptophan into a different position. However, there were no significant fluorescence intensity changes in 14:1 PC in the presence of PIP_2_, suggesting that in a more tilted TM configuration, the JM residues are less sensitive to electrostatic interactions with PIP_2_ (Fig. 3A). There were no significant spectral maxima changes for either 14:1 PC or 22:1 PC upon the addition of PIP_2_ (Fig. S5).

When we performed FRET experiments to determine the effect of PIP_2_ on the JM region, we observed differences in FRET efficiency across a range of acceptor concentrations, as before (Fig. S6). Fig. 3B shows that in thick bilayers, the presence of PIP_2_ decreased FRET by roughly half. This indicates that PIP_2_ increases the distance between the JM tryptophan and the DNS-labeled lipid headgroups. By contrast, in thin bilayers, PIP_2_ induced no significant changes in FRET efficiency (Fig. 3B). These results agree with the Trp fluorescence intensity changes caused by PIP_2_ observed in thick bilayers (Fig. 3A).

### PIP_2_ drives increased oligomerization of TMJM only in thick membranes

Since changes in oligomerization accompany changes in EphA2 signaling in cells (26, 39), we next determined if the interactions observed between the JM residues and PIP_2_ influenced the oligomerization of the peptide. Specifically, we used the single-molecule TIRF approach to examine the effect of PIP_2_ on the self-assembly of TMJM, in thin and thick SMALPs. We determined via TEM that the addition PIP_2_ did not alter SMALP size (Fig. S2). Fig. 4A shows that PIP_2_ increased the amount of dimeric TMJM peptide in 22:1 PC. The increased dimerization was reversed in the presence of Ca^2+^, indicating that this effect is due to an electrostatic interaction between the polybasic JM and the anionic PIP_2_ headgroup. While Ca^2+^ can destabilize SMALPs, this effect is observed only at higher concentrations (>2 mM) than we employed (40). In contrast, PIP_2_ did not affect dimerization in 14:1 PC (Fig. 4B), nor do the counts for larger oligomers demonstrate sensitivity to PIP_2_ (Fig. S4). These data agree with the tryptophan fluorescence and FRET data, which indicate that TMJM is sensitive to PIP_2_ only in thick bilayers. Our data suggest that PIP_2_ has a specific effect, which is limited to the conformation TMJM adopts in thick bilayers. Further, this effect is likely due to JM-PIP_2_ electrostatic interactions.

**Figure 4.**
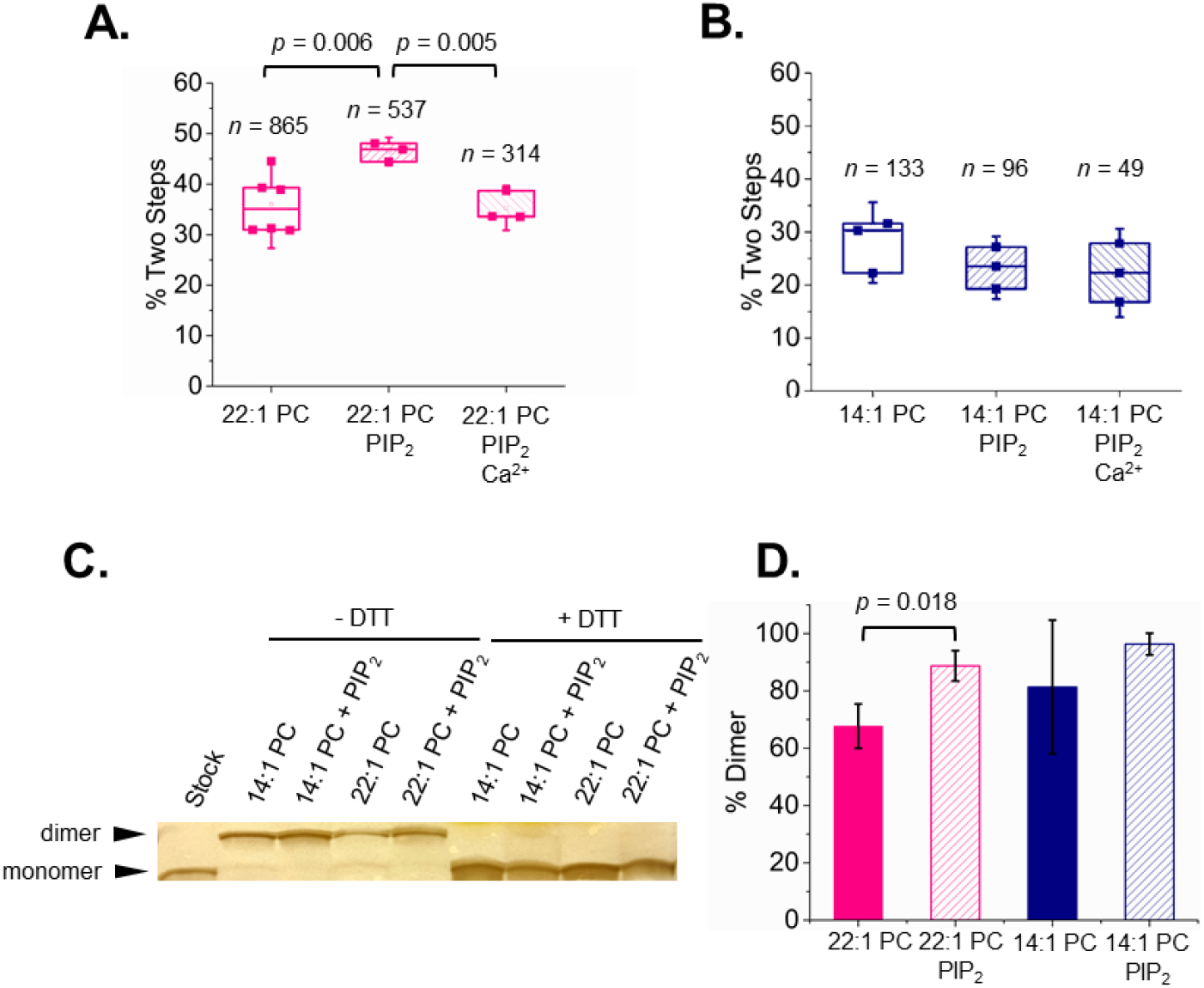
PIP_2_ promotes self-assembly of TMJM in thick bilayers. **A** and **B.** Percentage of Alexa Fluor 488 labeled TMJM peptide in two-step photobleaching traces in 22:1 PC and 14:1 SMALPs, respectively, determined by single-molecule studies. The effect of the presence of 3% PIP_2_ and 1 mM Ca^2+^ are investigated. Data are from 3-6 independent experiments. *n* is number traces counted for each. *p*-values are from student’s t-tests with significance determined after Benjamini-Hochberg procedure. **C.** Representative SDS-PAGE (full gel can be seen in Fig. S9) of unlabeled TMJM in 14:1 PC and 22:1 PC liposomes in the presence and absence of 3% PIP_2_. Monomer and disulfide-mediated dimers can be seen in non-reducing conditions. Addition of 5 mM DTT eliminates the disulfide-mediated dimer band. **D.** Quantification of 3 independent SDS-PAGE experiments as shown in C. Bands in each lane were quantified in ImageJ and percent of dimer was calculated. Bars are means ± S.D., *p*-value is from a student’s t-test. All data are from experiments at a lipid to peptide ratio of 300:1.

To further investigate the effects of JM interactions on the self-assembly of TMJM, we performed SDS-PAGE experiments where dimerization was investigated by disulfide cross-linking. We reasoned that if PIP_2_-dependent changes in self-assembly are promoted by lipid-JM interactions, this effect would be observable as differences in band sizes on a protein gel. Instead of measuring TMD-TMD interactions, as in the single-molecule experiments, we instead examined JM-JM interactions. To do this, TMJM containing a free Cys residue at the JM end was reconstituted in liposomes of 14:1 PC and 22:1 PC in the presence and absence of 3% PIP_2_. The peptide stock remained monomeric in our basal conditions (Fig. 4C, first lane). In the absence of a reducing agent, we observed two bands corresponding to monomer and disulfide-mediated dimer in all lipid conditions (Fig. 4C). The relative percentage of monomer and dimer was determined for each lipid condition. We observed that in 22:1 PC + PIP_2_ vesicles, the percentage of the disulfide-mediated dimer was higher than in 22:1 PC alone (Fig. 4D), in agreement with the single-molecule data in SMALPs. In 14:1 PC, no effect of PIP_2_ was observed.

### PS alters JM environment but not dimerization in thick membranes

Given the effects of PIP_2_ on TMJM, we wondered if phosphatidylserine (PS), an anionic lipid abundant at the plasma membrane, would also exert similar effects on the JM residues and dimerization of TMJM. The net charge of PS is −1, while PIP_2_ has on average a charge of ~−3 (41). To achieve a similar net charge in our model membranes, we used 22:1 PC with 10% PS (compared with 3% PIP_2_). This value is similar to PS levels found in the plasma membrane of eukaryotic cells (42). We tested for changes in tryptophan fluorescence and oligomerization in 22:1 PC, where PIP_2_-dependent changes were observed previously. A significant increase in fluorescence intensity was observed in the presence of PS (Fig. 5A). However, unlike the PIP_2_ experiments, this increase was not fully reversed in the presence of saturating amounts of Ca^2+^, suggesting differences in the effect of the two anionic lipids. Furthermore, single-molecule experiments showed that the presence of PS did not promote dimerization (Fig. 5B). These data suggest that PS alters the environment of the JM residues like PIP_2_, but without simultaneously driving significant changes in dimerization.

**Figure 5.**
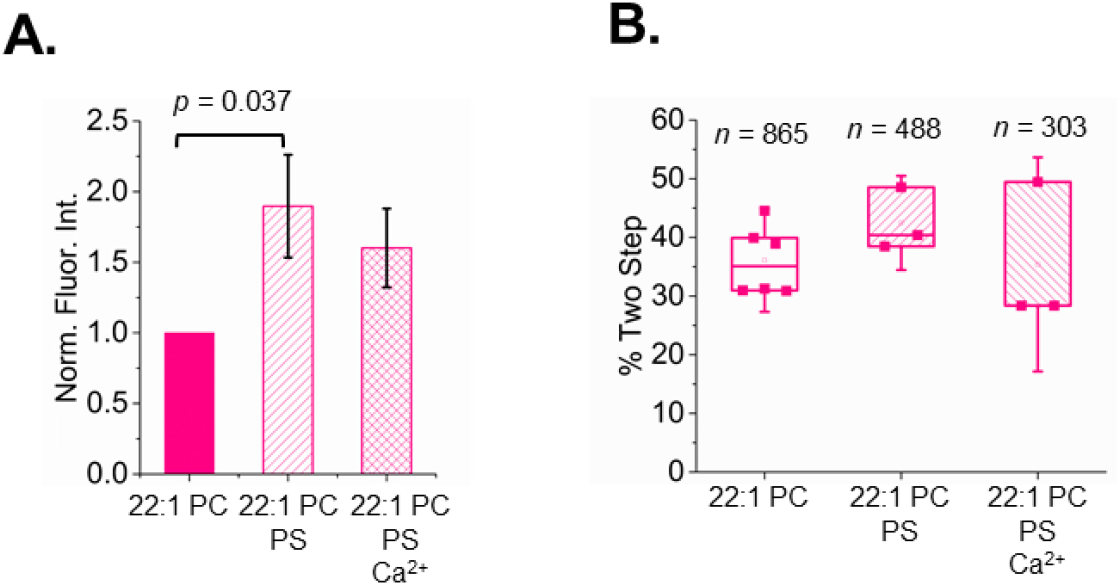
PS interactions with TMJM in thick bilayers. **A.** Normalized fluorescence intensities from emission spectra of TMJM in 22:1 PC liposomes with 10% POPS and 5 mM Ca^2+^. Bars are means ± SD, *n* = 3. *p*-value was determined by Mann-Whitney U test. **B.** Percentages of dimeric TMJM from SM-photobleaching experiments in 22:1 PC examining effects of 10% POPS and 1 mM Ca^2+^. Data are from 3-6 independent experiments. *n* is number traces counted for each. No statistically significant differences were found.

## Discussion

We have developed a reconstitution system that stabilizes the membrane region of the EphA2 receptor in two different conformations. OCD experiments performed in 14:1 PC bilayers indicated that the TMJM peptide adopted a highly tilted TM orientation (Fig. 1B), while in 22:1 PC bilayers the α-helix aligns more perpendicular to the bilayer plane (43). It is theoretically possible to use OCD to calculate specific helical tilt angles (44), but we could not accurately carry out this approximation due to uncertainties in peptide density in the supported bilayers employed. However, there is an intriguing qualitative agreement between our data and the two TM conformations reported for EphA2 (28, 29). The ligand-dependent dimer crosses the membrane in a highly tilted state (45°), while the ligand-independent dimer has a small crossing angle (15°) (28, 29). We propose that the two conformations found in our experimental conditions might correspond to the structure of the membrane region of EphA2 in the two different activation states: in 22:1 PC dimerization would occur with almost parallel helices, as expected for the ligand-independent dimer, and in 14:1 PC, a high-crossing angle dimer would correspond to the conformation induced after ephrinA1 binding. Our data indicate that conformational selection can be accomplished using thin and thick bilayers by taking advantage of the strong propensity of TMDs to avoid hydrophobic mismatch (32).

To ensure that the TMJM engages in biologically relevant dimerization we employed two complementary methods. After finding artifacts in ensemble FRET experiments performed in liposomes, we endeavored to develop a new single-molecule approach that uses SMALPs, which have been shown to maintain native membrane structures (Fig. 2) (36, 37). We also performed a crosslinking assay in vesicles of the same lipid compositions as the SMALPs. While this second method does not measure dimerization *per se*, it is expected to report on a related event, the proximity of the JM cysteines. With this limitation in mind, it is possible that the methods do not give an exact measurement of the amount of dimer in equilibrium, but an adequate estimation to make comparisons between different lipid environments. Not surprisingly, the two methods yielded different levels of total dimerization of TMJM. However, they agree in reporting comparable levels of the dimer found in both 14:1 PC and 22:1 PC (compare Fig. 2C and 4C and D). We conclude from the dimerization and OCD data that our thin and thick bilayer systems promote two different helical orientations of TMJM, and that in both cases the peptide can form a dimer.

The JM segment of EphA2 is functionally important, as it contains residues Y588 and Y594, which can be phosphorylated by the kinase domain of EphA2. This event triggers the release of the receptor from the auto-inhibited state (17). Hedger *et al*. examined the interaction of basic JM residues of 58 RTKs with anionic lipid headgroups via MD studies (30). They concluded that JM residues closest to the TM establish significant contacts with PIP_2_. Specifically for EphA2, their simulations predicted that the HRRRK region of the JM contributed the most to contacts with PIP_2_. Based on this observation, we included the HRRRK residues in the TMJM peptide. Similar observations have been made in simulations of the JM and kinase domain of EphA2 in bilayers containing PIP_2_ (45). However, these JM-PIP_2_ interactions have never been experimentally demonstrated.

Using the tryptophan near the C-terminus as a sensor, we were able to assess the JM environment in different lipids. Typically, the burial of tryptophan in a membrane results in a blue shift of the spectral maximum and a concurrent increase in fluorescence intensity (38). The change in bilayer thickness significantly affected the tryptophan fluorescence intensity, but no accompanying shift in spectral maximum was observed. This uncoupling of intensity and spectral maximum could be due to adjacent residues that quench tryptophan fluorescence. Specifically, the neighboring cysteine residue can engage in excited-state electron transfer with tryptophans (46). Further, it has been shown that tryptophan fluorescence spectra are sensitive to nearby charged residues (47). This led us to conclude that the local environment of the tryptophan is different in thin and thick bilayers, not because of differences in membrane burial but due to changes in relative orientation or proximity to neighboring residues. When PIP_2_ was added to thick bilayers, the tryptophan fluorescence increased significantly and was reversed upon the addition of Ca^2+^. This observation indicates that an electrostatic interaction occurs between the polybasic JM residues and the anionic PIP_2_ headgroups. We did not observe fluorescence changes with PIP_2_ in thin bilayers, indicating that the peptide is not sensitive to the charged lipids in this alternate conformation. To explain the increase in tryptophan fluorescence intensity and the decrease in FRET observed only in 22:1 PC membranes in the presence of PIP_2_ we have developed two possible explanations (Fig. 6). First, it is possible the JM becomes more buried into the core of the membrane moving the Trp away from the headgroups. Alternatively, PIP_2_ may cluster around the JM residues and this crowding pushes the DNS-PE headgroups away from the tryptophan laterally in the plane of the bilayer.

**Figure 6.**
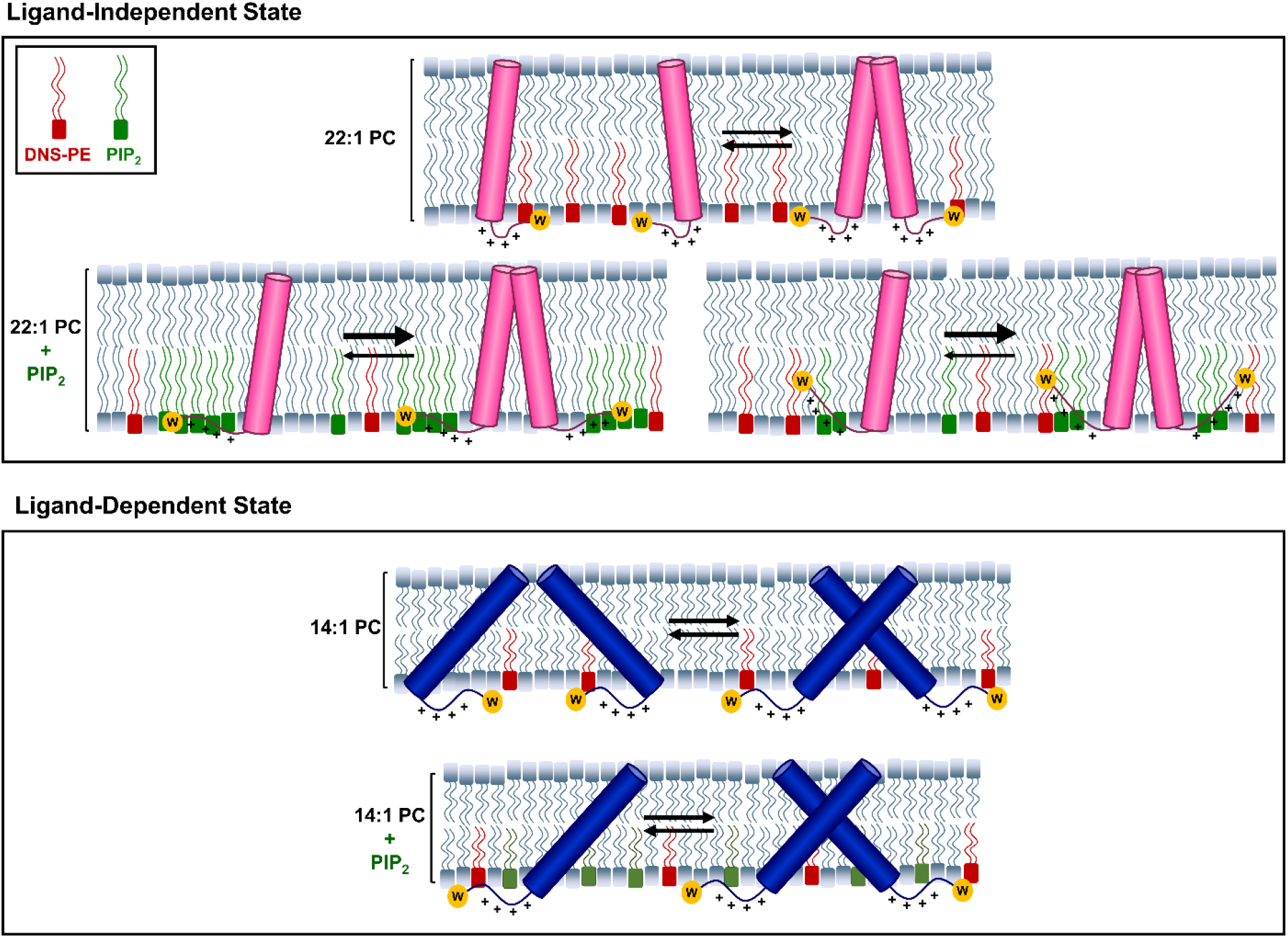
Model TMJM showing different effect of PIP_2_ on two TMJM configurations. **(top)** In the ligand-independent signaling configuration TMJM exists in a monomer-dimer equilibrium in the absence of PIP_2_. Charge-charge repulsion of the JMs must be overcome for efficient dimerization. In the presence of PIP_2_ the JM-lipid association changes by either clustering of PIP_2_ around the JM or burial of the JM. This shields the positive charges promoting dimerization. **(bottom)** In the ligand-independent signaling configuration, due to the tilt of the TM, charge repulsion of the JMs is not present. Neither JM environment nor dimerization is altered by PIP_2_.

In single-molecule experiments, PIP_2_ promoted TMJM dimerization in thick bilayers. Disulfide crosslinking experiments supported that PIP_2_ promotes self-assembly in 22:1 PC bilayers. In thin bilayers, no PIP_2_-dependent changes in disulfide-mediated dimerization were observed, also in agreement with the single-molecule data. Based on previous work (28, 29) and our data, we propose a model in which the EphA2 TMD and JM could synergize. In the ligand-independent state, EphA2 dimerization is mediated by the heptad repeat motif resulting in an upright TM orientation. In this configuration, the JM residues are tightly associated with the inner leaflet of the cell membrane via electrostatic interactions with PIP_2_. This interaction prevents electrostatic repulsion of the JM segments to make dimerization more energetically favorable. (Fig. 6, top). Without PIP_2_, EphA2 dimerization would be less favorable due to the charge-charge repulsion of the JMs. Taken together, our results suggest that interaction of PIP_2_ with the JM segments promotes TMD-TMD dimerization only when helices are roughly parallel.

On the other hand, in the ligand-dependent state, the TMD dimers would rotate to the glycine zipper dimerization interface and open to create a larger interhelical crossing angle. With the larger crossing angle of the dimer, charge-charge repulsion due to the JMs is minimized, and thus PIP_2_ would not promote dimerization. A relief of charge repulsion during ligand binding has been observed for the growth hormone receptor, where extracellular JM residues engage in charge-charge repulsion which is overcome by ligand binding and allows the TMD dimer to rotate and open to a larger inter-helical crossing angle (48).

Tryptophan fluorescence data with PS suggest that, as with PIP_2_, electrostatic interactions are enough to alter the environment of the JM region. However, unlike PIP_2_, the interaction with PS is not fully reversed upon addition of saturating amounts of Ca^2+^. This could be due to differences in binding sites and stoichiometry of the two lipids with Ca^2+^ ions (49, 50). Interestingly, PS did not promote dimerization, and the tryptophan fluorescence changes were larger. This leads us to conclude that JM interactions with PS do alter the environment of the JM but without promoting dimerization. In this case, the charge density of PS may not be large enough to overcome the charge-charge repulsion caused by the JM polybasic stretch. It has been reported that PS engages in contacts with RTK JM residues to a lesser degree than PIP_2_, which may explain why we see an uncoupling of the effect of PS on JM environment and oligomerization (30).

Our biophysical data suggest that PIP_2_ might play a direct role in modulating EphA2 signaling by stabilizing ligand-independent dimers and holding the phosphorylatable JM tyrosine residues at the plasma membrane. One study proposed that the ligand-independent oncogenic signaling is caused primarily by monomeric EphA2 (26). By promoting dimerization of the ligand-independent conformation, PIP_2_ would be reducing oncogenic signaling. However, dimerization via PIP_2_ in the unliganded state can be potentially opposed by the SAM domains which are known to inhibit oligomerization (57). PIP_2_-JM interactions have been demonstrated experimentally for several receptors including the epidermal growth factor receptor (EGFR), fibroblast growth factor receptor (FGFR), and tropomyosin receptor kinase A (TrkA) (51–54). It is believed that these electrostatic interactions serve to sequester phosphorylatable JM residues, rendering them inaccessible to the kinase domain, prior to ligand binding (51, 55). This effect is likely paired with the kinase domains also binding with the membrane, which has been shown via MD for EphA2, and experimentally for EGFR (55, 56).

We speculate that if ephrinA1 binds EphA2 in the ligand-independent dimeric conformation, the receptor would undergo rearrangements including TM rotation and opening of the crossing angle. Dimerization via the glycine zipper interface would not be promoted by PIP_2_, but by interactions between other parts of the full-length protein that oligomerize such as the cysteine rich domains and the ligand binding domains, which interact through two different interfaces upon ligand-binding (27, 39, 58). The glycine zipper TMD dimer may also be further stabilized by interactions with other proteins, which may contribute to the formation of large signaling clusters. For example, it is believed that interactions between SAM domains and dimers of SHIP2 may form large linear arrays (59, 60).

Our findings provide new insights that suggest that PIP_2_, and maybe other phosphorylated inositol lipids, could directly act as a conditional ligand of EphA2, causing Akt-independent modulation of the ligand-independent conformation of the receptor. We are currently testing the hypothesis that the lipid environment specifically regulates EphA2 signaling in cells.

## Experimental Procedures

### Liposome Preparation

Lipids were purchased from Avanti Polar Lipids, Alabaster, AL. 14:1 PC (1,2-dimyristoleoyl-*sn*-glycero-3-phosphocholine), 22:1-PC (1,2-dierucoyl-*sn*-glycero-3-phosphocholine), PIP_2_ (L-α-phosphatidylinositol-4,5-bisphosphate (Brain, Porcine)), 18:1 dansyl-PE (1,2-dioleoyl-*sn*-glycero-3-phosphoethanolamine-*N*-(5-dimethylamino-1-naphthalenesulfonyl), POPC (1-palmitoyl-2-oleoyl-glycero-3-phosphocholine), biotin-PE (1-oleoyl-2-(12-biotinyl(aminododecanoyl))-*sn*-glycero-3 - phosphoethanolamine), and PIP_2_ Bodipy FL (Echelon Biosciences, Salt Lake City, UT) stocks were prepared in chloroform. Aliquots of lipids were dried under argon gas and then placed in a vacuum overnight. Unless otherwise noted lipid films were resuspended with Buffer A: 19.3 mM HEPES (4-(2-hydroxyethyl)-1-piperazineethanesulfonic acid) buffer (pH 7.5), 1 mM EGTA (ethylene glycol-bis(β-aminoethyl ether)-*N,N,N’,N’*-tetraacetic acid), and 5 mM DTT (dithiothreitol). Large unilamellar vesicles (LUVs) were formed by extrusion with a Mini-Extruder (Avanti Polar Lipids, Alabaster, AL) through a 100 nm pore size membrane (Whatman, United Kingdom).

### Peptide Conjugation

The TMJM peptide was synthesized by F-moc chemistry by ThermoFisher (Waltham, MA), and purity (>95%) was assessed by MALDI-TOF and HPLC. The cysteine residue of TMJM was conjugated to Alexa Fluor 488 (Molecular Probes, Eugene OR) or Cyanine5 (Cy5) (Fluoroprobes, Scottsdale AZ), using a C5 maleimide moiety. The reaction was carried out by adding a molar excess (peptide:dye of 1:1.1 moles) of dye dissolved in 100 mM sodium phosphate pH 7.6 to a peptide stock in 2,2,2-trifluoroethanol (TFE). Unreacted dye was removed by HPLC, by injecting the TFE mixture onto a semi-preparative Agilent Zorbax 300 SB-C18 column on an Agilent 1200 HPLC system (Santa Clara, CA). The gradient of water + 0.05% trifluoroacetic acid (TFA) to acetonitrile + 0.05% TFA was 30 min from 0% - 100% acetonitrile. The conjugated peptide eluted around 95% acetonitrile. The collected fractions from HPLC were frozen and lyophilized. The dry conjugated peptide was resuspended in hexafluoroisopropanol (HFIP).

### Oriented Circular Dichroism

A stock of TMJM was prepared in TFE. Aliquots of 2.23 x 10^-7^ moles of 14:1 PC or 22:1 PC were dried under argon, then vacuum desiccated for at least 2 hours. The appropriate amount peptide stock (for a 50:1 or 300:1 lipid to peptide molar ratio) was added, dried with argon and finally dried under vacuum at least 2 hours. The lipid-peptide film was resuspended with 400 μl TFE and 150 μl spread on each of two circular quartz slides (Hellma Analytics, Germany). To allow for even solvent evaporation, the slides were placed in a fume hood overnight and further dried under vacuum for at least 6 hours to ensure complete evaporation of the TFE. The samples were hydrated under argon with 150 μl per slide of Buffer A overnight in 96% relative humidity, to obtain supported bilayers. Excess buffer was removed, and the hydrated slides were assembled in an OCD cell, with an inner cavity filled with saturated K2SO4 to keep the bilayers humidified. The OCD spectra were recorded on a Jasco J-815 spectropolarimeter at room temperature. For each sample eight 45° rotations of the cell were averaged. Appropriate lipid backgrounds were subtracted in all cases.

### Tryptophan Fluorescence

13 mm glass culture tubes (Fisher Scientific) were cleaned with piranha (75% H2SO4, 25% H2O2) solution for 3 minutes, creating a hydrophilic surface to promote efficient removal of peptide. Appropriate amounts of 100% 22:1 PC/14:1 PC or 3 mol% PIP_2_ and 97 mol% 22:1 PC/14:1 PC stocks were added to the cleaned tubes and dried under argon. Next, the lipids were dried under vacuum for at least 2 hours before the appropriate amount of peptide stock was added to achieve a lipid to peptide molar ratio of 300:1 and subsequently dried under vacuum overnight. Films were resuspended in Buffer A for an initial peptide concentration of 4 μM and extruded. To maximize peptide recovery, resuspension was conducted in three stages. First, 50% of the buffer volume was added to the tube then vortexed for 45 sec. This buffer was removed then the procedure was repeated twice with 25% of the final buffer volume. Equivalent lipid blanks were also prepared. To ensure that amounts of lipids between blanks and proteoliposomes were equal, ammonium molybdate phosphate assays were performed to quantify lipids (61). If necessary, lipid blank concentrations were appropriately adjusted. LUVs were then diluted to 300 μM lipid and 1 μM peptide ± 5 mM CaCl2 (where indicated). Samples were incubated for a minimum of 1 hour at room temperature to allow calcium levels to equilibrate across the bilayer. Tryptophan fluorescence spectra were then collected on a Cary Eclipse Fluorescence Spectrophotometer (Agilent Scientific, Santa Clara, CA) using an excitation wavelength of 290 nm. Lipid blanks were subtracted in all cases.

### Trp-DNS FRET

Lipids and peptide were dried in piranha-cleaned glass tubes as described above. Films were resuspended as described above in Buffer A for an initial peptide concentration of 4 μM. Equivalent lipid blanks were also prepared. Liposomes and proteoliposomes containing 0% and 10% dansyl-PE were mixed in appropriate ratios and subjected to seven rounds of freeze thaw to achieve 0%, 0.25%, 0.5%, 1%, 2%, 3%, and 5% dansyl-PE, ± 1 μM peptide, and ± 5 mM CaCl2 final concentrations where indicated. Samples were incubated at room temperature for a minimum of 1 hour to allow calcium levels to equilibrate. FRET experiments were conducted on a Cary Eclipse Fluorescence Spectrophotometer (Agilent Scientific, Santa Clara, CA) using an excitation wavelength of 290 nm. FRET efficiencies (E) were calculated with the following equation:

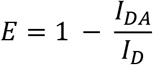

Where I_DA_ is the intensity of the donor in the presence of acceptor and I_D_ is the intensity of the donor only.

### Calcium Influx Assay

POPC vesicles were prepared by resuspending dried POPC with Buffer A, and 0.1 mM Indo-1 (1H-Indole-6-carboxylic acid, 2-[4-[bis-(carboxymethyl)amino]-3-[2-[2-(bis-carboxymethyl)amino-5-methylphenoxy]ethoxy]phenyl]-, pentapotassium salt). LUVs were formed via extrusion as described above. To separate encapsulated and free Indo-1, LUVs were subjected to size exclusion chromatography on a sephadex G25 PD-10 column (GE Life Sciences, Chicago, Il). The concentration of the encapsulated Indo-1 was estimated using fluorescence and known amounts of free Indo-1. Indo-1 containing LUVs were diluted to a final Indo-1 concentration of 0.05 μM in Buffer A and 5 mM CaCl2 was added. Calcium influx was observed in a Cytation5 plate reader (BioTek, Winooski, VT) as a shift in fluorescence maximum from 485 nm to 405 nm. Calcium influx saturated after 5 minutes. Free dye with 5 mM CaCl2 and encapsulated dye with 0.1% Triton X-100 were used as controls.

### SMALP preparation

For photobleaching experiments, peptide and lipid films were prepared by drying down 22:1 PC or 14:1 PC + 3% biotin-PE ± 3% PIP_2_ or 10% POPS from chloroform stocks. To this film TMJM conjugated with Alexa Fluor 488 in HFIP was added. For co-localization experiments, lipids and peptides were prepared the same way as in photobleaching experiments with PIP_2_ Bodipy FL and TMJM conjugated to Cy5. The amount of lipid was kept constant while the amount of peptide was adjusted for the specified lipid to peptide ratio (300:1, 100:1 or 50:1). The films were dried under Ar gas, then vacuum-desiccated for at least two hours. MLVs were then formed by resuspending in SMALP buffer (19.3 mM HEPES, 150 mM NaCl, 1 mM EGTA). MLVs were subjected to three rounds of freeze-thaw at −80 °C and 42 °C to ensure even mixing of the lipid components. A stock solution of 1-9 mg/mL of SMA 2000H (Polyscope, Geleen, The Netherlands) was diluted to 0.3 mg/mL and added to MLVs for a SMA final concentration of 0.075 mg/mL and lipid concentration of 100 μM. The MLV/SMA solution was then incubated overnight with shaking to allow SMALP formation.

### Single-molecule TIRF

Quartz microscope slides (G. Finkenbeiner Inc., Waltham MA) and coverslips were cleaned following the protocol of Chandradoss *et al*. 2014 (62). Clean slides and coverslips underwent animosalinization and pegylation following a procedure previously described (63). In short, slides were incubated with a solution of 93% methanol, 4.5% acetic acid and 2.5% 3-(triethoxysilyl)-propylamine (EMD Millipore Corp., Billerica, MA), rinsed with methanol and water and finally dried under a stream of nitrogen (63). A solution of m-PEG-SVA and biotin-PEG-SVA (Laysan Bio Inc., Arab, AL) was made by dissolving 20% w/v m-PEG and 1.25% biotin-PEG in filtered 100 mM NaHCO3 overnight. To assemble a flow chamber, slides were pre-drilled with holes and fitted with a coverslip using double sided tape and sealed with vacuum grease. Pegylated slides were incubated for 10 min with 0.2 mg/mL streptavidin, and then washed with SMALP buffer. SMALPs, containing a peptide concentration of 20-30 nM, were immobilized on the slides for 10 min and then rinsed to remove any nonspecific interactions. The rinse buffer was replaced with an oxygen scavenging system; 2.5 mM PCA and 250 nM rPCO (recombinant Protocatechuate 3,4-Dioxygenase; Oriental Yeast Co., Tokyo Japan) in SMALP buffer with 2 mM Trolox (64). Slides were imaged under a custom-based TIRF microscope and the emission intensities were collected on CCD camera (Andor Technology) with 100 ms integration time. A custom written software package (downloaded from https://physics.illinois.edu/cplc/software) was used to record movies and extract single-molecule traces using scripts written in IDL (Harris Geospatial Solutions, Inc) software (65). Single-molecule traces were assessed and analyzed using custom software written in Python and analyzed to determine the number of photobleaching steps. To prevent potential bias, the experimenter was blinded during analysis with a custom data-shuffling Python script.

### SDS-PAGE

Lipid-peptide films were prepared for SDS PAGE as described above with unlabeled TMJM peptide at a lipid to peptide ratio of 300:1. Dried lipid-peptide films were resuspended in 19.3 mM HEPES, 1 mM EGTA and shaken at room temperature for 3 hours to allow disulfide bond formation. To the MLVs, SDS buffer was added for a final SDS concentration of 150 mM. To this, sample buffer +/- DTT was added. Samples were then boiled for 5-10 minutes to ensure complete disruption of liposomes. To ensure the stock of peptide did not contain disulfide-mediated dimers, a sample of the TMJM stock was also prepared without lipid. This sample was resuspended in buffer containing 150 mM SDS and loaded with sample buffer without DTT. Samples were run on a 16% tricine gel and stained using a Peirce Silver Stain Kit (Thermo Scientific, Waltham MA). Bands were quantified in ImageJ using the Band Peak Quantification plugin.

### Transmission Electron Microscopy

SMALPs were prepared as described above to a final lipid concentration of 1 mM. 3% w/w SMA was added to both lipids before overnight equilibration. SMALPs of 14:1 PC were equilibrated with shaking at room temperature overnight, while 22:1 PC SMALPs were incubated at 60 °C for the same duration. SMALPS were imaged with negative staining TEM. Small aliquots of SMALPs were adsorbed on to glow discharged carbon-coated copper EM grids (Electron Microscopy Sciences, Hatfield, PA) for 120 s. Grids were washed twice with ddH2O for 15 s before negative staining with UranyLess (Electron Microscopy Sciences, Hatfield, PA) for 45 s. Excess liquid was removed from the grid with filter paper between steps. Grids were air-dried prior to examination on a JEOL JEM 1400-Flash TEM (JEOL USA, Peabody, MA) operating at 80 kV. SMALPs were measured in ImageJ.

### Statistical Analysis

All statistical comparisons were made in IBM SPSS v25 software (Armonk, New York USA). Where only two means were compared, student’s t-tests were used. Where more than two means were compared one-way analysis of variance (ANOVA) were conducted followed by post-hoc comparisons, with Tukey HSD tests where data were homoscedastic, and Dunnett’s t-test where data were heteroscedastic. All *p*-values reflect an α = 0.05. Where no *p*-values are shown, the difference is not significant.

## Data Availability

Data are available upon reasonable request.

## Acknowledgments

This work was supported by NIH grant R01GM120642 (FNB). We are thankful to Adam Smith and Blake Mertz for ideas on the project, and to Ms. Jennifer Schuster, Dr. Haden Scott, Ms. Alyssa Ward, and Mr. Nicolas Moore for their thoughtful comments on the manuscript.

## Abbreviations

TM: transmembrane
JM: juxtamembrane
RTK: receptor tyrosine kinase
ALS: amyotrophic lateral sclerosis
MD: molecular dynamics
HR: heptad repeat
GZ: glycine zipper
SMALPs: styrene maleic acid lipid particles
TEM: transmission electron microscopy
TIRF: total internal reflection fluorescence
OCD: oriented circular dichroism
FRET: Förster Resonance Energy Transfer
ANOVA: analysis of variance
PS: phosphatidylserine
DTT: dithiothreitol
MALDI-TOF: matrix assisted laser desorption ionization time of flight
HPLC: high performance liquid chromatography

