## Supporting Information for "PIP_2_ promotes conformation-specific dimerization of the EphA2 membrane region"

### $PIP_2$ promotes conformation-specific dimerization of the EphA2 transmembrane domain

**Supplementary Figures:**

Supplemental Figure 1. CD of TMJM at 50:1 and OCD at 300:1

Supplemental Figure 2. TEM images and size distributions of SMALPs

Supplemental Figure 3. Controls for immobilization of SMALPs on slides via biotin-streptavidin linkage and SMALP composition.

Supplemental Figure 4. Percent larger oligomers of TMJM in SMALPs.

Supplemental Figure 5. Tryptophan fluorescence spectra and spectral maxima.

Supplemental Figure 6. Tryptophan-DNS FRET spectra and efficiencies at different acceptor concentrations.

Supplemental Figure 7. OCD of TMJM in 22:1 PC and 14:1 PC with $PIP_2$.

Supplemental Figure 8. Peptide recovery and calcium influx controls.

Supplemental Figure 9. Full SDS-PAGE gel of TMJM crosslinking.

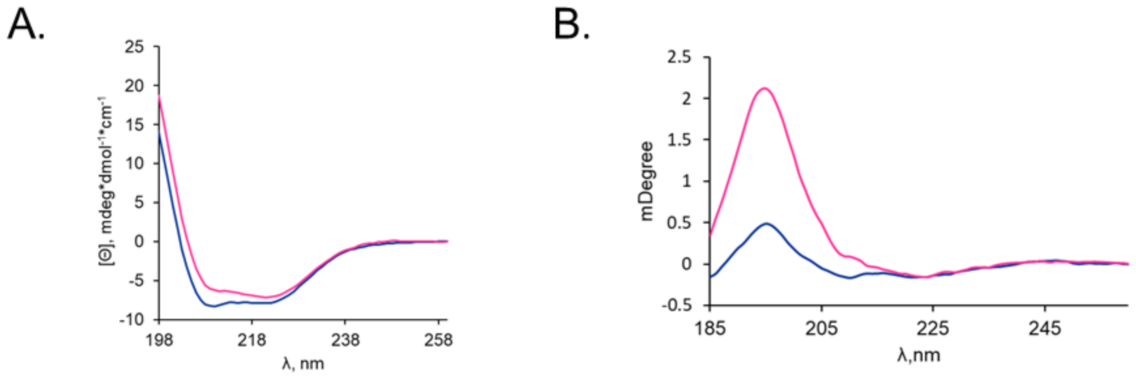


**Supplemental Figure 1**. **A.** CD of TMJM in 22:1 PC (fuchsia) and 14:1 PC (navy) liposomes at a lipid to peptide ratio of 50:1 showing that the secondary structure in both lipids is α-helical. **B.** OCD of TMJM in 14:1 PC (navy) and 22:1 PC (fuchsia) at a lipid to peptide ratio of 300:1.

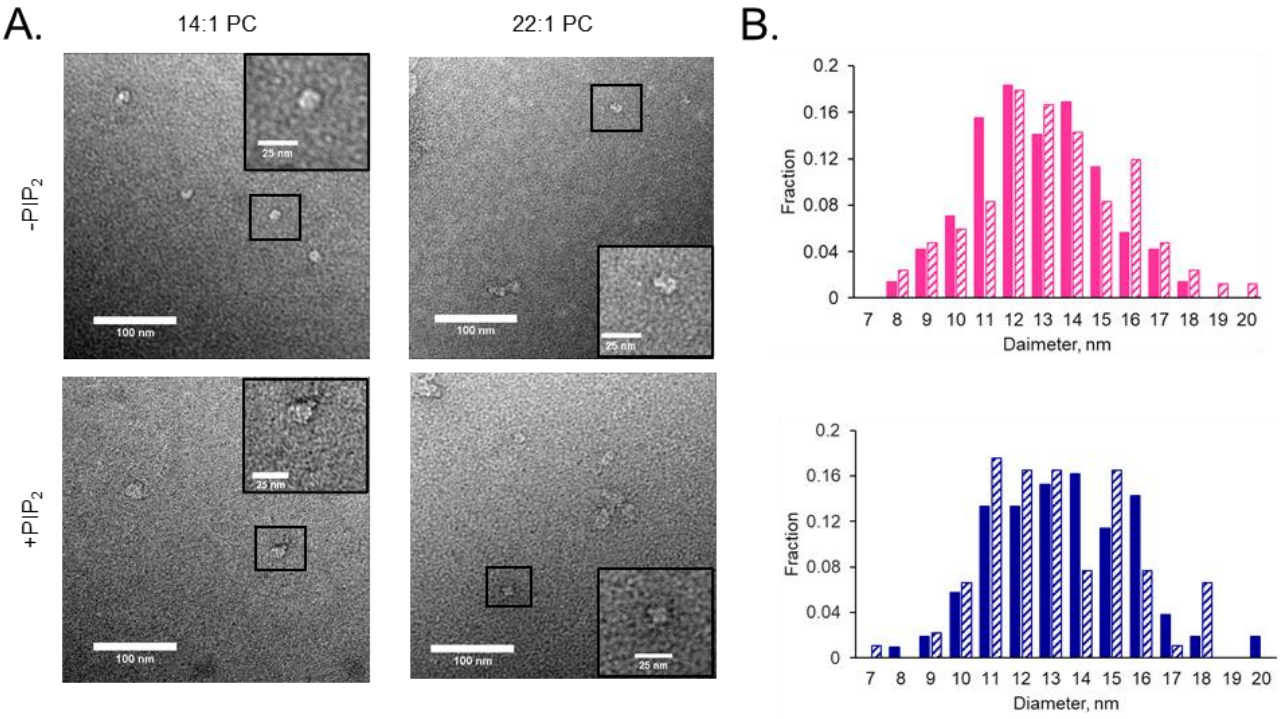


**Supplemental Figure 2. A.** Representative TEM images of SMALPs comprised of 22:1 and 14:1 PC ± 3% $PIP_2$. Scale bars in large images are 100 nm. Scale bars in insets are 25 nm. **B.** Histograms of SMALP diameters. Top: 22:1 PC (solid bars) and 22:1 PC + $PIP_2$ (cross hatch). Bottom: 14:1 PC (solid bars) and 14:1 PC + $PIP_2$ (cross hatch). Data are from 3-4 independent SMALP preparations for each lipid composition. 80-100 SMALPs were measured for each lipid composition.

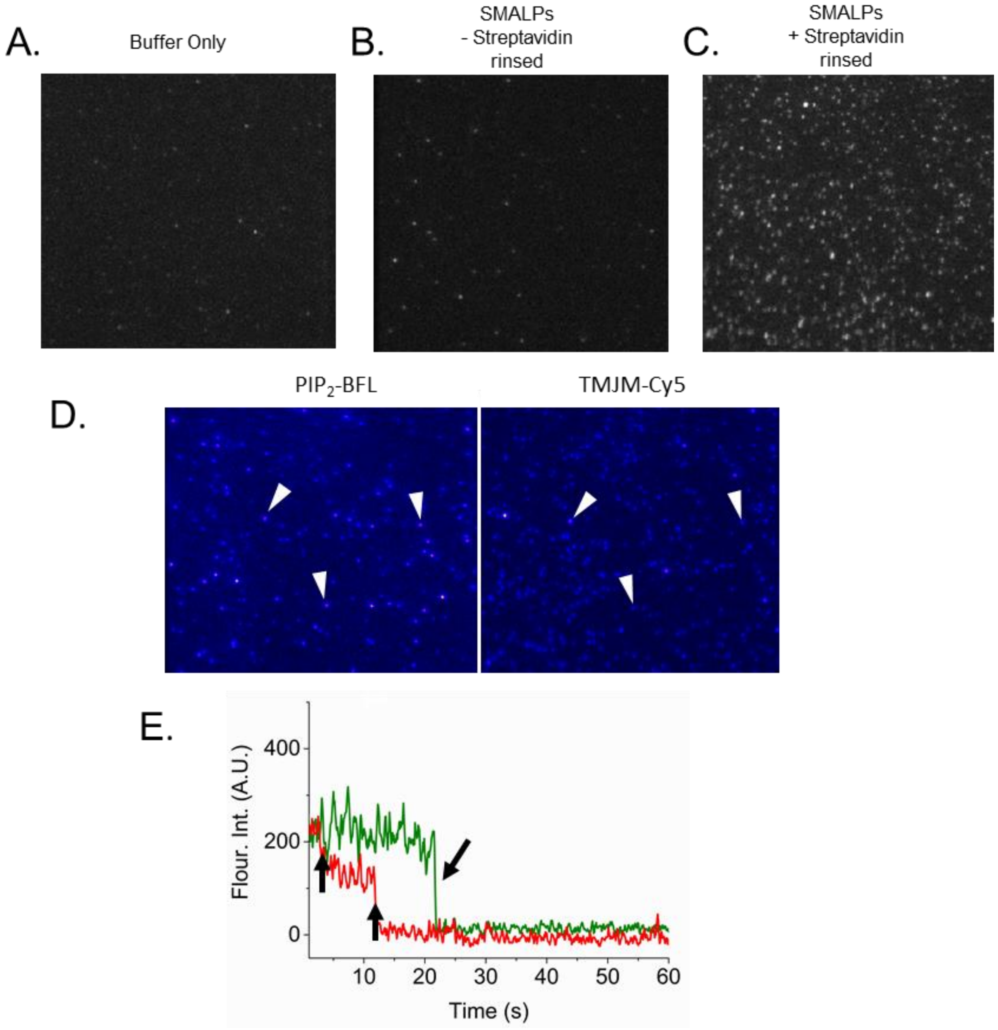


**Supplemental Figure 3. Controls for immobilization of SMALPs on slides via biotin-streptavidin linkage and SMALP composition. A. Control** biotinylated slide with buffer only. **B.** A biotinylated slide without streptavidin was incubated for 10 minutes with SMALPs containing 3% biotin-PE and TMJM Alexa Fluor 488 and rinsed. Images show no non-specific immobilized SMALPs. **C.** A biotinylated slide was incubated with 0.2 mg/mL streptavidin for 10 min followed by incubation with SMALPs containing biotin-PE and TMJM Alexa488 then rinsed. Image shows immobilized SMALPs. **D.** Co-localization of SMALPs with $PIP_2$ Bodipy FL (left) and TMJM Cy5 (right) simultaneously excited. White arrows highlight examples of SMALPs containing both $PIP_2$ and TMJM. **E.** Representative fluorescent trace from co-localization data in panel D showing TMJM Cy5 (red) with two photobleaching steps (black arrows) and fluorescence from $PIP_2$ Bodipy FL (green) showing one photobleaching step (black arrow).

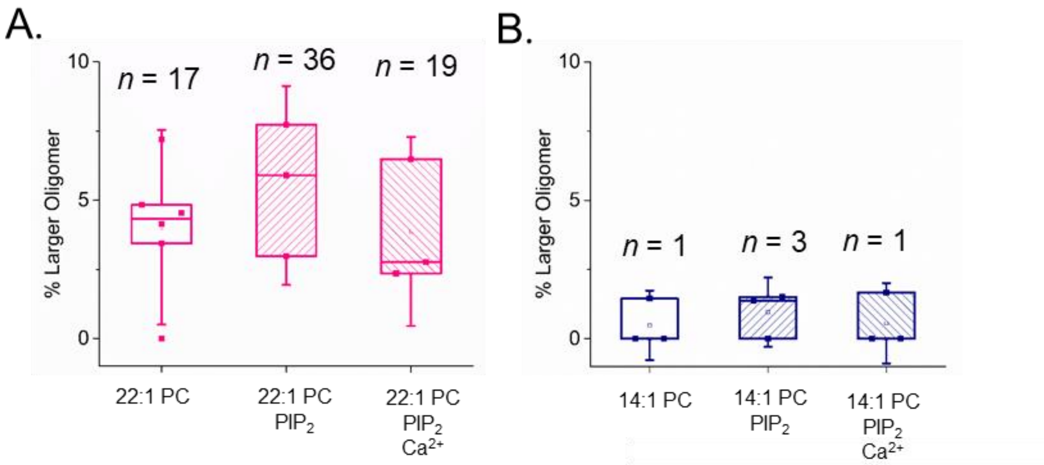


**Supplemental Figure 4. A.** and **B.** Percentage of peptide in larger oligomers in 22:1 PC and 14:1 PC SMALPs with and without 3% $PIP_2$ and 5 mM $Ca^{2+}$ via SM-photobleaching experiments. Data are from 3-6 independent experiments. *n* = number of traces counted with 3 or more steps.

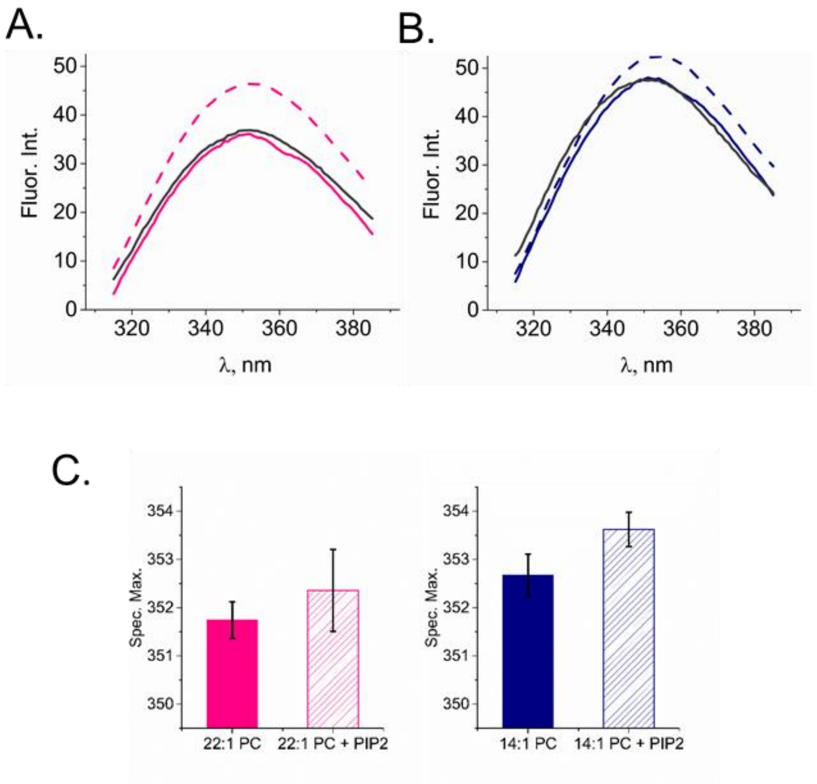


**Supplementary Figure 5. A.** Trp emission spectra in 22:1 PC liposomes (solid fuchsia line), with 3% $PIP_2$ (dashed line) and 3% $PIP_2$ with $Ca^{2+}$ (gray line) (curves are averages of 3 independent experiments). **B.** Trp emission spectra in 14:1 PC liposomes (solid navy line), with 3% $PIP_2$ (dashed line) and 3% $PIP_2$ with $Ca^{2+}$ (gray line). Curves are averages of 3 independent experiments. **C.** Tryptophan fluorescence spectral max in 14:1 PC and 22:1 PC liposomes with and without 3% $PIP_2$. Bars are means ± S.D. from 3 independent experiments.

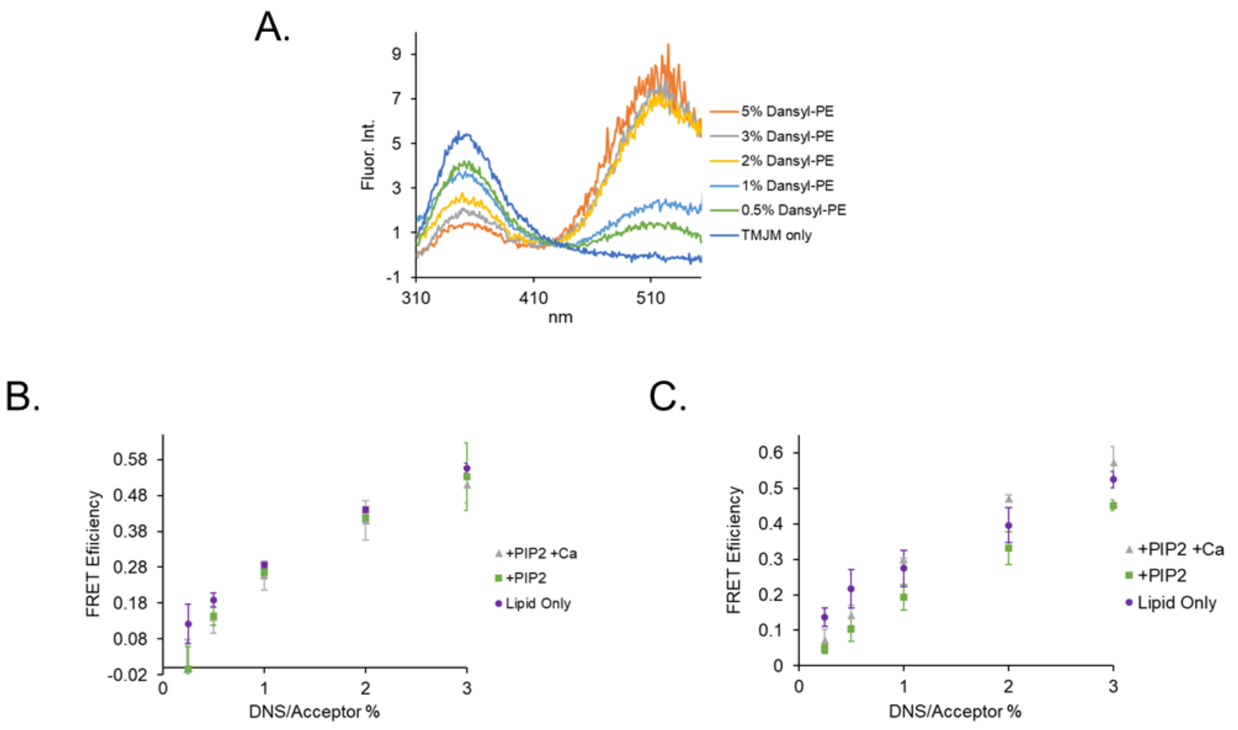


**Supplemental Figure 6. A**. Representative emission spectra of Trp and Dansyl showing saturating amounts of FRET at ~2% DNS-PE. **B.** FRET efficiencies of 1 µM TMJM in LUVs with 0-3% DNS-PE without $PIP_2$ or $Ca^{2+}$ (purple), with 3% $PIP_2$ (green), and with 3% $PIP_2$ and 5 mM $Ca^{2+}$ (grey) in 14:1 PC liposomes. **C.** FRET efficiencies of 1 µM TMJM in LUVs with 0-3% DNS-PE without $PIP_2$ or $Ca^{2+}$ (purple), with 3% $PIP_2$ (green), and with 3% $PIP_2$ and 5 mM $Ca^{2+}$ (grey) in 22:1 PC liposomes. Points are averages of 3 independent experiments ± S.D.

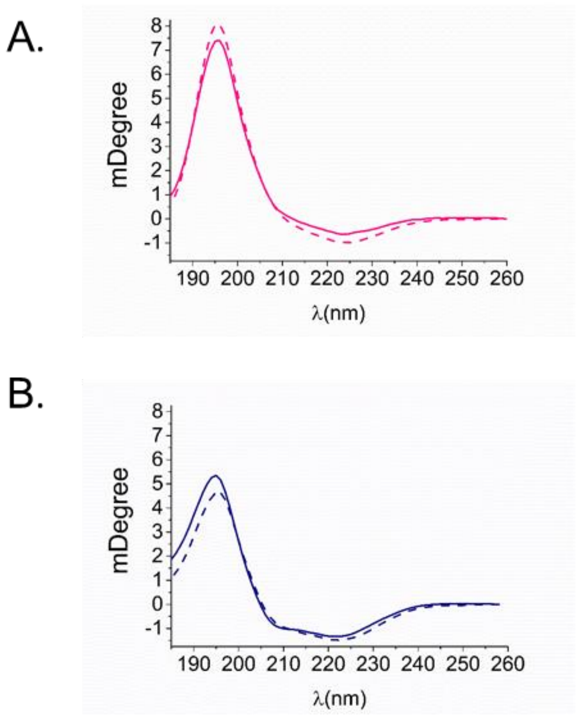


**Supplemental Figure 7. Helical tilt in thin and thick bilayers is preserved upon addition of $PIP_2$. A.** OCD spectra of TMJM in 22:1 PC with $PIP_2$ (dashed line) and without $PIP_2$ (solid line). Curves are averages of 3 independent experiments. **B.** OCD spectra of TMJM in 14:1 PC with $PIP_2$ (dashed line) and without $PIP_2$ (solid line). Curves are averages of 3 independent experiments.

A.

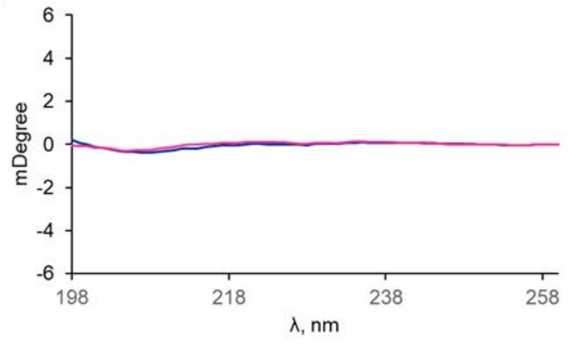


B.

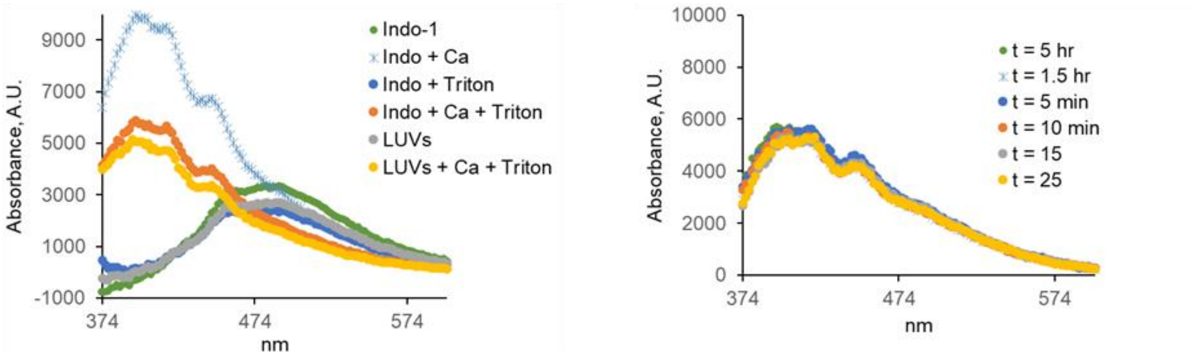


**Supplemental Figure 8. A.** Representative CD spectra of SDS-washed tubes indicating comparable levels of peptide were recovered for 22:1 PC (fuchsia) and 14:1 PC (navy) liposomes in fluorescence experiments. **B. (left)** Blue shift of encapsulated Indo-1 dye spectral maximum is observed after addition of 5 mM $Ca^{2+}$. **(right)** Calcium influx assays showing 5 mM $Ca^{2+}$ crosses the membrane in saturating amounts within 25 minutes.

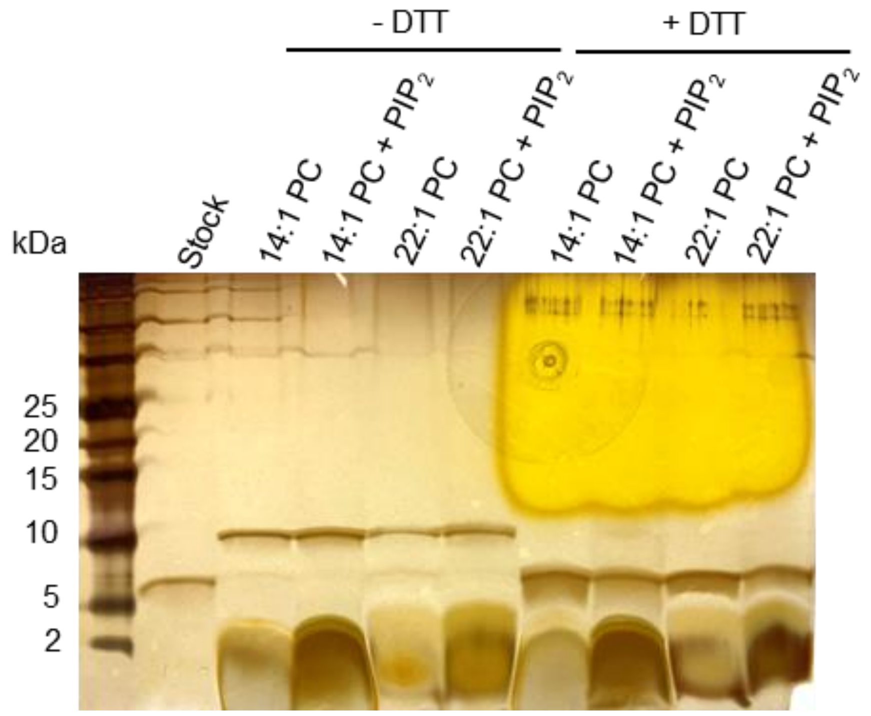


**Supplemental Figure 9.** Representative SDS PAGE gel of TMJM crosslinking as shown in Fig. 4C. Smears below 5 kDa are from lipids. Note: no smear is seen in first lane where no lipid was added. Yellow discoloration in upper-right portion of gel is from DTT, as the faint very high bands observed.
